# miR-16 Suppresses Growth of Rhabdoid Tumor Cells

**DOI:** 10.1101/219709

**Authors:** Emily Kunce Stroup, Yunku Yeu, Albert Budhipramono, Tae Hyun Hwang, Dinesh Rakheja, Anat Erdreich-Epstein, Theodore W. Laetsch, James F. Amatruda, Kenneth S. Chen

## Abstract

**Background:** Rhabdoid tumor is a highly aggressive pediatric cancer characterized by biallelic loss and/or mutation of *SMARCB1*. Outcomes remain poor, and there are no established ways to target the tumorigenic pathways driven by SMARCB1 inactivation. *SMARCB1* loss leads to an increase in cyclin D transcription.

**Procedure:** We characterized the cell line WT-CLS1, which has been described previously as Wilms tumor, by whole-exome sequencing, RNA-seq, and xenograft histology. We measured the effect of microRNA overexpression on WT-CLS1, BT-12, and CHLA-06-ATRT.

**Results:** We found that WT-CLS1 demonstrates the histological, mutational, and transcriptional hallmarks of rhabdoid tumor. Because the microRNAs let-7 and miR-16 can target cyclin D genes, we next overexpressed each of these microRNAs in WT-CLS1. We found that miR-16 reduced cell accumulation. This was accompanied by a decrease in proliferation markers and an increase in apoptosis markers. These results were replicated in the BT-12 and CHLA-06-ATRT cell lines.

**Conclusions:** The loss-of-function *SMARCB1* mutation found in WT-CLS1, in conjunction with immunohistochemical and gene expression analysis, warrants reclassification of this cell line as rhabdoid tumor. Proliferation of WT-CLS1 and other rhabdoid tumor cell lines is significantly abrogated by miR-16 overexpression. Further studies are necessary to gain insight into the potential for miR-16 to be used as a novel therapeutic in rhabdoid tumor.

**Abbreviations:**

(ATRT): atypical teratoid/rhabdoid tumor
(miRNA): microRNA
(dox): doxycycline
(OD595): optical density at 595 nm
(CDK4): cyclin-dependent kinase 4

## INTRODUCTION

Rhabdoid tumor is a rare, highly aggressive cancer which usually occurs before the age of four. It can arise from the brain, the kidney, or other extracranial sites. In the brain, it is known as “atypical teratoid/rhabdoid tumor” (ATRT)^1^. Despite the use of intensive chemotherapy regimens, including high-dose chemotherapy with autologous stem cell rescue, prognosis remains poor regardless of anatomic site^1–4^. Among renal tumors, rhabdoid tumor was recognized as a separate entity from Wilms tumor in 1981^5^. Rhabdoid tumors of the kidney are historically treated on high-risk Wilms tumor protocols, though prognosis remains poorer for patients with rhabdoid tumor compared to Wilms tumor^6–8^.

Rhabdoid tumors are marked molecularly by biallelic inactivation of *SMARCB1* (also known as SNF5, BAF47, or INI1)^9,10^. Aside from *SMARCB1* loss, rhabdoid tumors have remarkably simple mutational profiles^11–13^. These molecular features are shared by ATRT and extracranial rhabdoid tumors alike. SMARCB1 is a core subunit of the multiprotein SWI/SNF chromatin remodeling complex, which is frequently mutated in cancer ^14^. SMARCB1 mediates SWI/SNF binding at differentiation-regulated enhancers, where it guides distinct epigenetic and gene expression signatures^11,12,15,16^. Loss of SMARCB1 leads to dysregulation of a variety of cellular pathways, most notably activation of cyclin D1 and cyclin-dependent kinase 4 (CDK4)^17–20^. Specifically, SMARCB1 inhibits expression of D-type cyclins, and *SMARCB1* loss leads to overexpression of these cyclins. Based on these findings, efforts to therapeutically target SMARCB1-deficient rhabdoid tumors aim to specifically block cyclin D activity^21,22^.

In cancer and other diseases, RNA-based therapeutic approaches are emerging as a novel pharmacological approach towards difficult therapeutic targets. One approach uses microRNAs (miRNAs), which are small ubiquitous noncoding RNAs that regulate the expression of target genes through base-pairing with their 3’ untranslated regions^23^. They are dysregulated in many cancers; for instance, about 20% of Wilms tumors harbor mutations in the enzymes responsible for microRNA processing^24–27^. In particular, let-7 mimics have been shown to impair growth of rhabdoid tumor cells *in vitro* To our knowledge, no other microRNAs have been studied as potential therapeutic strategies in rhabdoid tumor.

Here we show that the cell line WT-CLS1, which was previously described as Wilms tumor, displays the histological, gene expression, and mutational characteristics of rhabdoid tumor. Additionally, we show that miR-16, which targets all three members of the cyclin D family, potently inhibits growth of WT-CLS1 and other rhabdoid tumor cell lines. This may be a novel therapeutic strategy for rhabdoid tumor.

## METHODS

### Cell Culture

The WT-CLS1 cell line was obtained from Cell Line Services (Eppelheim, Germany) and cultured in Iscove’s Modified Dulbecco’s Medium (Gibco, Waltham, MA) supplemented with 10% FBS and 1x antibiotic-antimycotic (Gibco, Waltham, MA). Its identity was confirmed by short tandem repeat profiling. BT-12 was obtained from the Children’s Oncology Group Cell and Xenograft Repository, and CHLA-06-ATRT was generated as previously described^29,30^. Both of these ATRT lines were maintained in Iscove’s Modified Dulbecco’s Medium with 20% FBS and 1x antibiotic-antimycotic.

### Subcutaneous Xenograft and Histological Staining

Xenografts were established in 8-week-old NOD-*scid* IL2RYγ^null^ mice by subcutaneous injection of 5 million cells suspended in a 1:1 mixture of Matrigel (Corning, Corning, NY) and serum-free media. After 2-3 months, these tumors were processed and stained for immunohistochemistry or with hematoxylin and eosin. Immunohistochemistry for INI-1 was performed using the standard immunoperoxidase method on an automated immunostainer (Dako Omnis, Agilent, Santa Clara, CA, USA). After heat-induced epitope retrieval at high pH, a mouse monoclonal primary antibody for SMARCB1 (clone MRQ-27, Cell Marque, Rocklin, CA, USA) was used at 1:25 dilution.

### Sequencing and Genomic Analysis

DNA was prepared using the DNeasy Blood & Tissue Kit (Qiagen, Hilden, Germany) and submitted to DNA Link (San Diego, CA) for whole exome sequencing. Exome capture was performed using Agilent SureSelect (Agilent Technologies, Santa Clara, CA) and sequenced on the Hiseq 2000 platform (Illumina, San Diego, CA) using 100bp paired-end sequencing. Reads were aligned to the hg19 reference genome using BWA (v0.7.10). Duplicates were marked with Picard tools (v1.119), and Genome Analysis Toolkit (GATK) v3.5.0 HaplotypeCaller was used to call variants. Known variants were then filtered out using published databases from the 1000 Genomes Project (v2015Aug), the Single Nucleotide Polymorphism database (dbSNP, v147), Exome Aggregation Consortium (ExAC, v0.3), and NHLBI Exome Sequencing Project (SI-v2).

RNA was prepared using the mirVana miRNA Isolation Kit (ThermoFisher Scientific, Waltham, MA) and submitted to DNA Link (San Diego, CA) for RNA sequencing. cDNA libraries were constructed by using KAPA Stranded RNA-Seq Kit with RiboErase (HMR) (Kapa Biosystems, Inc). Amplified cDNA was validated and quantified on an Agilent Bioanalyzer with the High Sensitivity DNA chip. The purified libraries were normalized, pooled, denatured, and diluted, before single-end 75 bp sequencing on NextSeq500 (Illumina, San Diego, CA). From each sample, we obtained about 25 million reads. Reads were aligned to the hg19 reference genome using STAR (v2.4.2a)^31^. Ensembl database (GRCh37.82) was used to guide the spliced transcript alignment. Read counts, in transcripts per million (TPM), were calculated using the FeatureCounts algorithm in the Subread aligner package (v1.5.1)^32^ Principal components analysis was performed in Python (v2.7) on Wilms tumor, rhabdoid tumor, and clear cell sarcoma of the kidney samples from the TARGET project (https://ocg.cancer.gov/programs/target/data-matrix). RNA-seq from G401 was downloaded from the ENCODE project^33^.

### Transduction of pSLIK Construct

The genomic regions surrounding hsa-let-7a-1 and hsa-miR-16-2 were cloned from human DNA into the XhoI site of the pSLIK-Neo-TGmiR-Gb2 plasmid (Addgene 25746)^34^ using the primers listed in Supplementary Table 1. Lentivirus was generated from these plasmids in HEK293T cells, and WT-CLS1 cells infected with these lentiviruses were selected in G418 (ThermoFisher Scientific, Waltham, MA). miRNA expression was induced by treating cells with doxycycline at 1 μg/ml. RNA was extracted using Trizol, and qPCR was performed using the Taqman Assay Kit for hsa-miR-16-5p (Assay #000391) and U6 (Assay #001973) (ThermoFisher Scientific, Waltham, MA).

Cells were plated at a density of 3×10^3^ cells/well in a 24-well plate in four replicates. Plates were fixed and stained using 6 mM crystal violet in 20% methanol. The stain was solubilized in 10% acetic acid and absorbance was measured at 595 nm. The media (with or without doxycycline) was replaced every 2 days.

### Western Blot and qPCR for miR-16 Targets

Immunoblots were performed on SDS-PAGE using PCNA and cleaved PARP antibodies (#13110 and #9542, respectively; Cell Signaling Technology, Danvers, MA), diluted 1:1000. qPCR was performed using iTaq Universal SYBR Green Supermix (Bio-Rad, Hercules, CA) with the primers listed in Supplementary Table 1.

### Transfection of miRNA Mimics

Control and miR-16 miRIDIAN miRNA mimics (Dharmacon, Lafayette, CO) were transfected into BT-12 and CHLA-06-ATRT cells using Lipofectamine RNAiMAX (ThermoFisher Scientific, Waltham, MA). Cells were seeded at 2.5×10^5^ cells/well in a 12-well plate and transfected using 2.5 μL RNAiMAX incubated with 30 pmol of each miRNA mimic in each well. Twenty-four hours post-transfection, these cells were evenly split into multiple 24-well plates. Each day four wells per condition were stained with crystal violet to measure cell density.

## RESULTS

### WT-CLS1 represents rhabdoid tumor

To characterize the WT-CLS1 cell line, we generated subcutaneous xenografts in immunocompromised mice. Hematoxylin and eosin-stained sections of formalin fixed and paraffin embedded xenograft tumors derived from WT-CLS1 cell lines were examined (Fig. 1A). They showed sheets of medium-sized cells with ovoid to irregular nuclei, vesicular chromatin, prominent eosinophilic nucleoli, well-defined nuclear membranes, and a moderate amount of pale eosinophilic cytoplasm that in many cells is aggregated into an inclusion-like body. This appearance resembled that of rhabdoid tumors. Accordingly, we performed immunohistochemistry, which demonstrated a loss of nuclear staining for SMARCB1 in the tumor cells.

**Figure 1.**
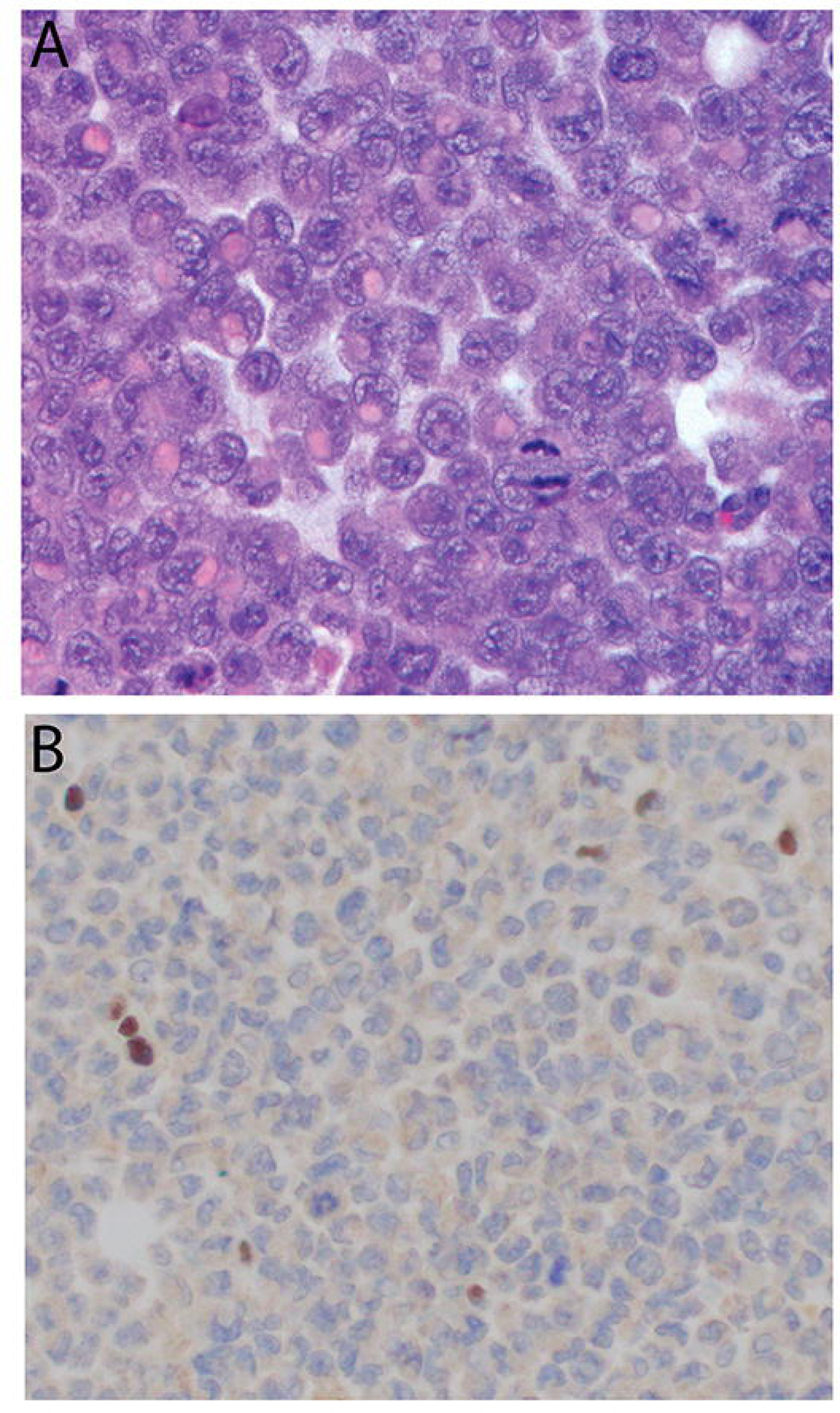
WT-CLS1 xenograft histology recapitulates rhabdoid tumor. (A) A photomicrograph of xenograft tumors derived from WT-CLS1 cell line shows the classic histologic features of a rhabdoid tumor with prominent nucleoli and inclusion-like eosinophilic cytoplasm (hematoxylin and eosin, 400 X original magnification). (B) Immunostain for SMARCB1 shows loss of nuclear staining in the tumor cells, while a few lymphocytes in the background show a normal pattern of nuclear staining (immunoperoxidase, 400 X original magnification).

We next characterized WT-CLS1 using whole-exome sequencing. This analysis revealed a nonsense mutation (Q318*) in all 40 of the reads spanning this genomic location, which produces a premature stop codon (Fig. 2A). We confirmed this finding with Sanger sequencing (Fig. 2B). It has previously been shown that WT-CLS1 cells have single-copy loss of chromosome 22, which contains the *SMARCB1* locus^35^. Thus, our results show that the second allele of *SMARCB1* is inactivated by nonsense mutation.

Having found that WT-CLS1 cells carry the morphological, immunohistochemical, and mutational signatures of rhabdoid tumor, we next studied its gene expression profile using RNA-seq (Supplementary Table S2). We used principal components analysis to compare its gene expression profile to that of other kidney tumors and cell lines (Fig. 2C). We accessed publicly available RNA-seq data from the TARGET project from Wilms tumors^25^, rhabdoid tumors^12^, and clear cell sarcomas of the kidney^36^, the three most common kidney tumors in pediatrics. We also included RNA-seq data from WiT49, a known Wilms tumor cell line, and publicly available RNA-seq data from G401, a known rhabdoid tumor cell line^37^. Projection of these data along the principal components axes showed that WT-CLS1 more closely resembled rhabdoid tumor than Wilms tumor.

**Figure 2.**
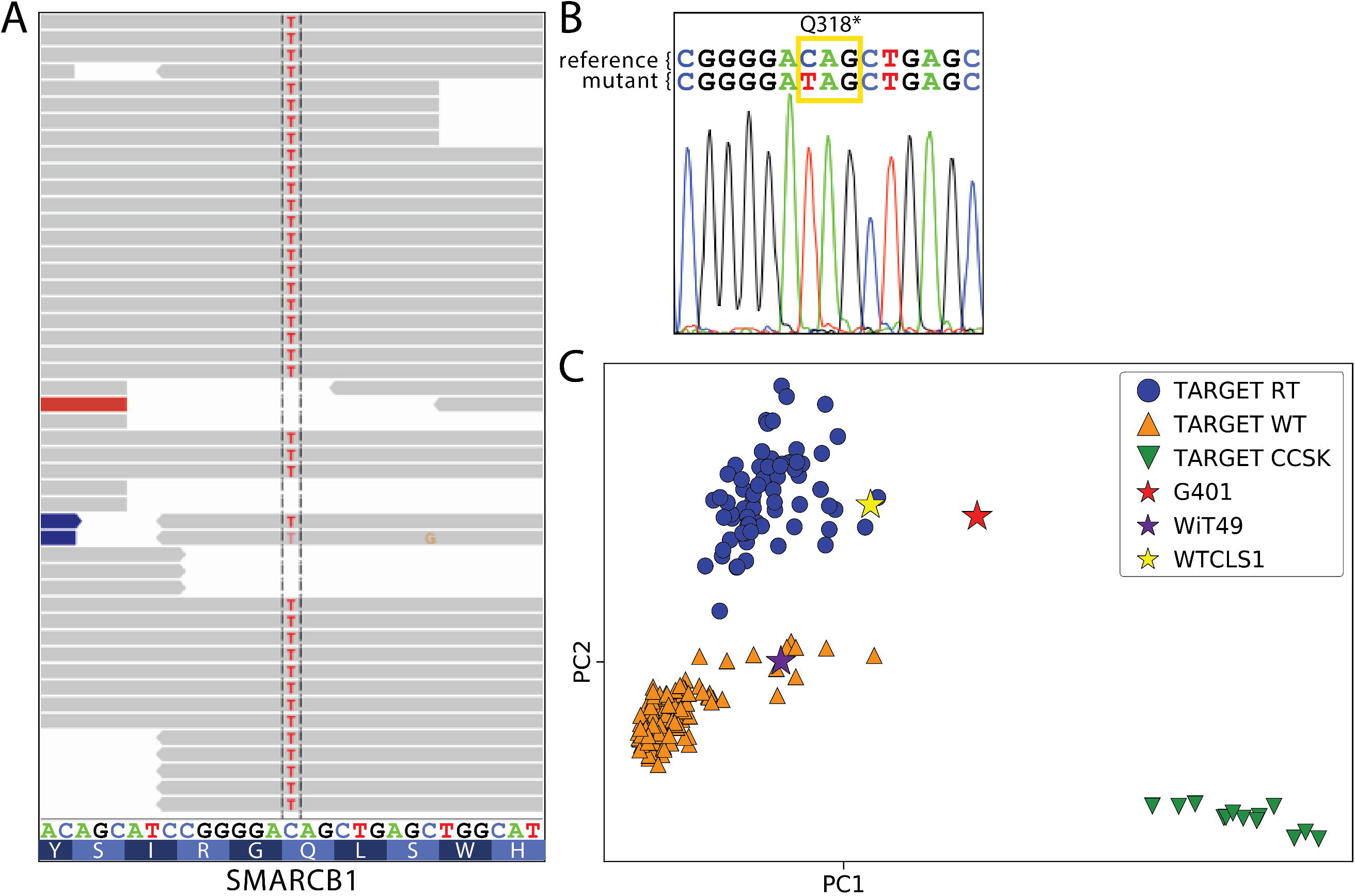
Next-generation sequencing analyses confirm features of rhabdoid tumor in WT-CLS1. (A) Whole exome sequencing of WT-CLS1 reveals a premature stop codon at amino acid 318 (p.Q318*) in 40 of 40 reads. (B) Sanger sequencing confirms truncating SMARCB1 mutation. (C) Principal components analysis of RNA-seq data for rhabdoid tumor (RT), Wilms tumor (WT), and clear cell sarcoma of the kidney (CCSK), compared to three cell lines (G401, WiT49, and WT-CLS1).

### miR-16 impairs WT-CLS1 proliferation

Having established that WT-CLS1 represents rhabdoid tumor, we next sought to identify novel therapeutic approaches in rhabdoid tumor. Specifically, we looked for microRNAs that could repress cyclin D genes. There are two microRNA families that are predicted to target all three cyclin D genes in a highly conserved manner based on TargetScan (v7.1)^38^: the let-7 and miR-16 families (Fig. 3A). These two microRNAs are well-characterized as tumor suppressors in other cancers, and they are both known to target a wide range of oncogenes^39,40^. A previous study suggested let-7 therapeutic replacement for rhabdoid tumor^28^. Thus, we decided to test whether each of these microRNAs could repress growth of WT-CLS1.

We used lentiviral vectors to establish stable sublines of WT-CLS1 which would express either let-7 or miR-16 upon doxycycline treatment. Strikingly, we found that cell density of WT-CLS1 was significantly decreased by miR-16 induction, but only mildly impaired by let-7 induction (Figs. 3B-3D). Next, we studied the molecular effects of miR-16 induction to understand its striking effect on cell accumulation. We confirmed by quantitative PCR that miR-16 induction leads to decreases in the expression of *CCND2* and *CCND3*, which are more highly expressed than *CCND1* in WT-CLS1 (Fig. 3E). Furthermore, we also used immunoblots to measure the effect of miR-16 on markers of proliferation and apoptosis. Induction of miR-16 led to both a decrease in proliferation (measured by proliferating cell nuclear antigen [PCNA]) and an increase in apoptosis (measured by poly-ADP-ribose polymerase [PARP] cleavage; Fig. 3F). These results suggested that miR-16 replacement may be a novel way to arrest growth of rhabdoid tumors.

**Figure 3.**
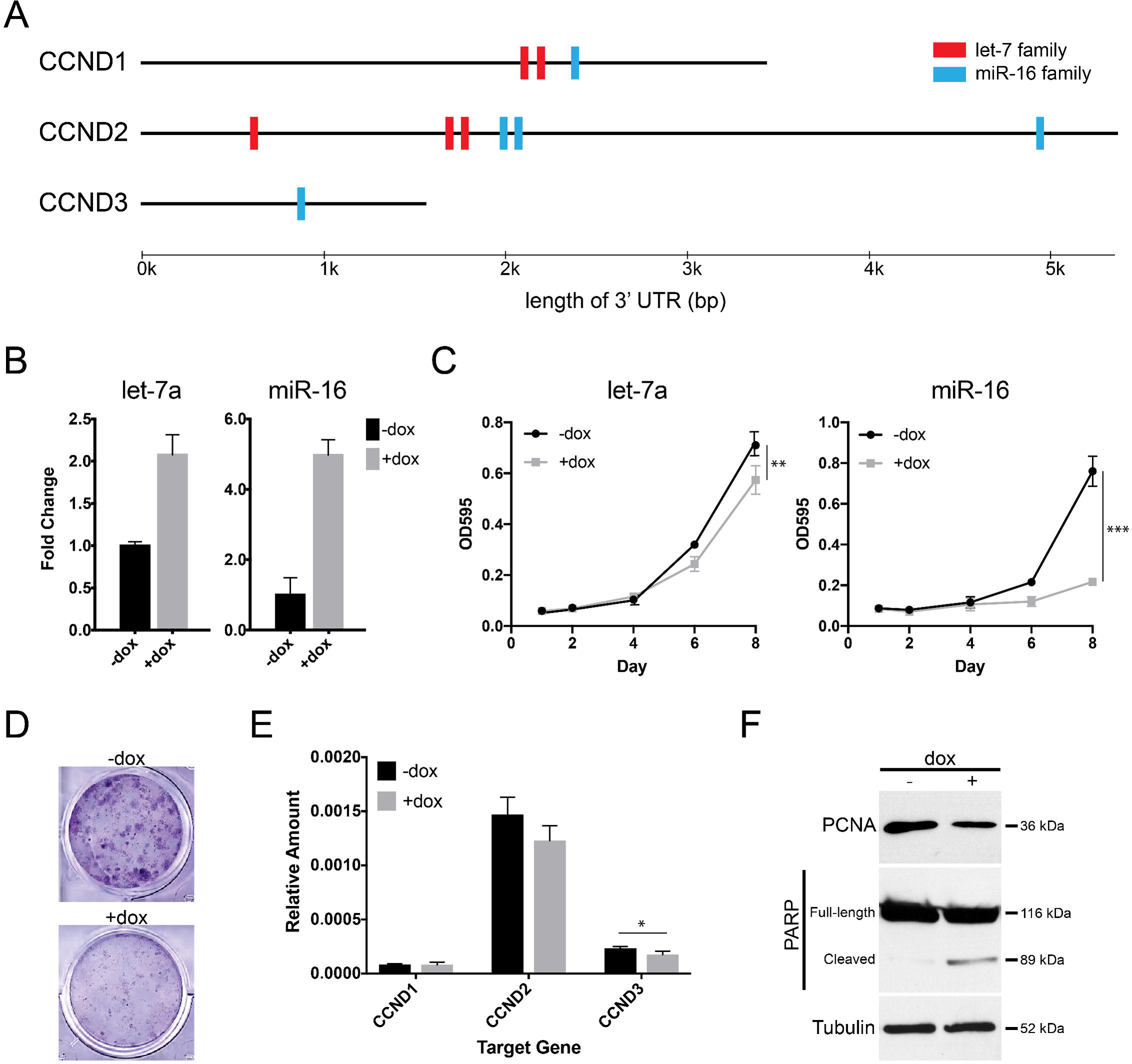
miR-16 impairs WT-CLS1 proliferation. (A) Highly conserved miRNA binding sites in the 3’ untranslated regions of cyclin D family gene, based on TargetScan v7.1. (B) WT-CLS1 cells lentivirally transduced to overexpress let-7a or miR-16 upon addition of doxycycline (dox). (C) Cell density of WT-CLS1 upon dox-induced let-7 or miR-16 expression, assayed by crystal violet. (D) Representative image of crystal violet-stained wells with and without doxycycline after 8 days. (E) qPCR for cyclins D1, D2, and D3 in setting of dox-induced miR-16 overexpression (* *p* < 0.05). (F) Immunoblots for PCNA and cleaved PARP show decreased proliferation and increased apoptosis following miR-16 induction.

### miR-16 also impairs proliferation of other rhabdoid tumor cell lines

To understand whether miR-16 growth inhibition also extends to other rhabdoid tumor contexts, we next examined the impact of miR-16 on the proliferation of atypical teratoid/rhabdoid tumor cell lines. We transfected two established ATRT cell lines, BT-12 and CHLA-06-ATRT, with miR-16 mimics (Fig. 4A). Similar to WT-CLS1, we found that miR-16 expression impaired cell accumulation when compared to a control mimic (Fig. 4B-4D). Again, we found that these cells also had a concomitant decrease in expression of D-type cyclins (Fig. 4E-4F). Thus, we concluded that miR-16 replacement can arrest the growth of a variety of rhabdoid tumor cell lines.

**Figure 4.**
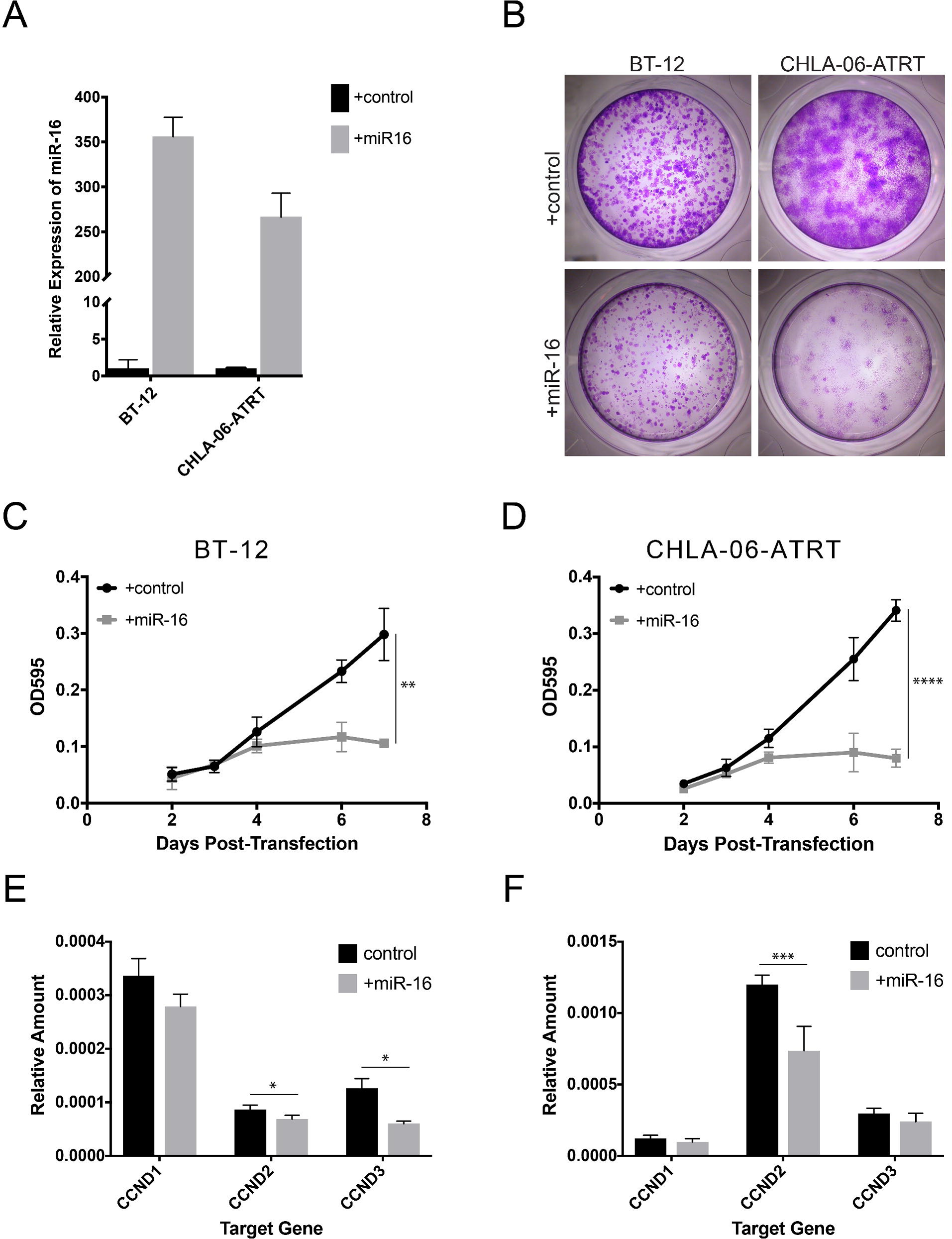
miR-16 impairs proliferation of BT-12 and CHLA-06-ATRT. (A) Transfection of miR-16 mimics in BT-12 and CHLA-06-ATRT. (B) Representative images of crystal violet-stained cells 8 days after transfection with control mimic or miR-16 mimic. (C,D) Cell density of BT-12 (C) and CHLA-06-ATRT (D) following miR-16 transfection. (* *p* < 0.05, ** *p* < 0.01, *** *p* < 0.001). (E,F) qPCR for cyclin D1, D2, and D3 in mimic-transfected cells.

## DISCUSSION

For years since its derivation, WT-CLS1 has been used as a model of Wilms tumor, but here we present evidence that it actually represents rhabdoid tumor. We show that WT-CLS1 has biallelic *SMARCB1* inactivation, at both the sequence and the immunohistochemical level. Additionally, its transcriptomic and morphological characteristics are consistent with rhabdoid tumor. Furthermore, we found that WT-CLS1 is sensitive to miR-16 overexpression. This is accompanied by knockdown of cyclin D genes, a known vulnerability for rhabdoid tumor. Lastly, we extend this finding to other rhabdoid tumor cell lines and show that they are also sensitive to miR-16 overexpression.

Wilms tumor cell lines have been difficult to isolate and culture long-term^41,42^. In fact, many cell lines which were originally described as Wilms tumor were later found to represent other cancers. For instance, soon after the distinction of rhabdoid tumor from Wilms tumor, the G401 cell line was found in fact to bear characteristics of rhabdoid tumor rather than Wilms tumor^37^. Likewise, SK-NEP-1, also previously described as a Wilms tumor, was found to in fact represent Ewing sarcoma due to presence of the EWS-FLI1 oncogenic fusion^43^. These findings forced a re-interpretation of many studies done using these cell lines, including the discovery of “WT2”, a purported second Wilms tumor suppressor gene, in G401^44^. Additionally, drugs that can inhibit G401 and SK-NEP-1 do not demonstrate efficacy against Wilms tumor^45–48^.

Several previous studies have found that features of WT-CLS1 were uncharacteristic of Wilms tumor without specifically identifying it as rhabdoid tumor. For example, it was previously shown that WT-CLS1 xenografts do not recapitulate the histological and immunohistochemical characteristics of human Wilms tumor^49^. Similarly, its gene expression pattern was unique from Wilms tumor cell lines, and it carried none of the common genomic aberrations common seen in Wilms tumor^50^. Lastly, unlike true Wilms tumor cell lines, WT-CLS1 showed a unique resistance to IGF1R inhibition^51^. Our re-characterization here explains these previously confusing results. On the other hand, our description of WT-CLS1 as a rhabdoid tumor may force re-interpretations of some conclusions drawn about Wilms tumor based on studies in WT-CLS1. For example, the importance of telomere shortening in WT-CLS1 may not be relevant to telomere lengths in human Wilms tumors^52^.

Although it has previously been shown that let-7 mimics impair rhabdoid tumor growth^28^, overexpression of let-7a in our tet-inducible system did not impair growth of WT-CLS1 as effectively as miR-16. This may reflect the high levels of let-7 at baseline in WT-CLS1, which limited the additional let-7a induction we could attain. miR-16 is a tumor suppressor in several other tumor types. In fact, miR-16 was the first tumor-suppressor microRNA discovered, when it was found that chronic lymphocytic leukemias commonly harbor deletion of the miR-15a/16-1 locus^53^.

Although we focus on its effects on the cyclin D family, miR-16 regulates genes in many oncogenic pathways, including cell cycle^40,54,55^, apoptosis^56^, invasiveness^39^, and chemoresistance^57^. In fact, a recent Phase I study of miR-16 mimics showed tolerable toxicities and some clinical responses in adults with malignant pleural mesothelioma, and ongoing studies are investigating its use in other cancers^58^. To our knowledge, this is the first study showing its potential use in rhabdoid tumor.

Our identification of WT-CLS1 as a rhabdoid tumor adds another tool for developing biological insights and novel therapies for this deadly cancer. Despite knowing the causative mutation for decades, outcomes for patients with rhabdoid tumor remain dismal. Further intensification of chemotherapy and radiation regimens has produced diminishing results and mounting toxicities. Studies on the oncogenicity of *SMARCB1* loss have led to several advances in our understanding of tumorigenesis in rhabdoid tumor. Our identification that miR-16 is suppressor of growth in rhabdoid tumor provides a novel alternative therapeutic strategy which bears further testing in future studies.

## ACKNOWLEDGMENTS

K.S.C.’s work was funded by an Alex’s Lemonade Stand Foundation Young Investigator grant and a Hyundai Hope on Wheels Young Investigator award.

## SUPPLEMENTARY FILES

**Supplementary Table S1**. Primers used in this study

**Supplementary Table S2**. RNA-seq table from WT-CLS1, in transcripts per million.

